# Exon Elongation Added Intrinsically Disordered Regions to the Encoded Proteins and Facilitated the Emergence of the Last Eukaryotic Common Ancestor

**DOI:** 10.1101/2022.04.03.486857

**Authors:** Satoshi Fukuchi, Tamotsu Noguchi, Hiroto Anbo, Keiichi Homma

## Abstract

Most prokaryotic proteins consist of a single structural domain (SD) with little intrinsically disordered regions (IDRs) that by themselves do not adopt stable structures, while the typical eukaryotic protein is comprised of multiple SDs and IDRs. How eukaryotic proteins evolved to differ from prokaryotic proteins has not been fully elucidated. Here, we found that the longer internal exons are, the more frequently they encode IDRs in eight eukaryotes including vertebrates, invertebrates, a fungus, and plants. Based on this observation, we propose the “small bang” model from the proteomic viewpoint: the protoeukaryotic genes had no introns and mostly encoded one SD each, but a majority of them subsequently divided into multiple exons (step 1). Many exons unconstrained by SDs elongated to encode IDRs (step 2). The elongated exons encoding IDRs frequently facilitated the acquisition of multiple SDs to make the last common ancestor of eukaryotes (step 3). One prediction of the model is that long internal exons are mostly unconstrained exons. Analytical results of the eight eukaryotes are consistent with this prediction. In support of the model, we identified cases of internal exons that elongated after the rat-mouse divergence and discovered that the expanded sections are mostly in unconstrained exons and preferentially encode IDRs. The model also predicts that SDs followed by long internal exons tend to have other SDs downstream. This prediction was also verified in all the eukaryotic species analysed. Our model accounts for the dichotomy between prokaryotic and eukaryotic proteins and proposes a selective advantage conferred by IDRs.

## Introduction

In contrast to the first eukaryotic common ancestor (FECA) that had no introns, the last eukaryotic common ancestor (LECA) probably had an intron-rich genome (Koonin 2006; Irimia and Roy 2014), but massive intron losses subsequently occurred in a few lineages of eukaryotes (Csuros et al. 2011). Alternative splicing produces multiple variants from one gene and can explain the variation in complexity among eukaryotes (Chen et al. 2014) and approximately 95% of multiexon human genes undergo alternative splicing (Pan et al. 2008). Introns, regarded as selectively disadvantageous, proliferated in the evolutionary process leading to the LECA probably by meiotic processes (Poole 2006). Recent research, however, uncovered a potential advantage of having introns: they attenuate the formation of DNA-RNA hybrids (R-loops) that are deleterious in transcription (Niu 2007). Prokaryotic proteins are comprised mostly of structural domains (SDs) and little intrinsically disordered regions (IDRs) that do not adopt stable three-dimensional structures by themselves, while on average ~33% of residues are in IDRs in eukaryotic proteins (Ward et al. 2004). In eukaryotic proteins IDRs are especially prevalent in transcription factors (Minezaki et al. 2006) and participate in the formation of biomolecuar condensates that regulate diverse cellular functions, particularly gene transcription (Fuxreiter and Vendruscolo 2021). IDRs can arise *de novo* by expansion of repetitive DNA sequences (Tompa 2003) and exonisation of introns (Kondrashov et al. 2003; Sorek 2007; Marquez et al. 2015). While prokaryotic proteins mostly consist of a single SD, eukaryotic proteins tend to have multiple SDs, which contribute to increased functional complexity (Koonin 2000; Koonin et al. 2002; Tordai et al. 2005). Domain accretion and the prevalence of IDRs in proteins are thus likely to be crucial in the evolution of complex cellular functions in eukaryotes. The exon theory proposed that exon shuffling by recombination within introns resulted in a gain in structural domains (Gilbert and Glynias 1993; Long et al 1995). However, the exon theory is not easily reconcilable with the observation that the boundaries of exons and the boundaries of structural domains of the encoded proteins do not align with high frequency, although they align more often than expected (Liu and Grigoriev 2004; Smithers et al. 2019).

We endeavored to propose a plausible evolutionary process that explains addition of IDRs and its effect on domain accretion in eukaryotic proteins. Although most human internal exons are between 50 to 200 nucleotides (nt) in length (Lander et al. 2001), 5% of them are longer than 1000 nt. These exons are evolutionarily conserved across mammals and are mostly expressed and translated (Bolisetty et al. 2012). The SRSF3 splicing factor promotes the inclusion of large exons enriched with C-nucleotides in vertebrates (Kawachi et al. 2021) and plays important roles in the occurrence and development of tumours (Xiong et al. 2022). Intriguingly, long exons in vertebrates frequently encode IDRs (Kawachi et al. 2021). We investigated if this tendency is observed in other eukaryotes and thus is likely to have been present in the LECA.

## Results

### The longer internal exons are, the more frequently they encode IDRs

We examined the internal exons of the eight model eukaryotes including three vertebrates (*Homo sapiens, Mus musculus* and *Rattus norvegicus*), two invertebrates (*Caenorhabditis elegans* and *Drosophila melanogaster*), one fungus (*Schizosaccharomyces pombe*), and two plants (*Arabidopsis thaliana* and *Oryza sativa* subsp. *japonica*). Internal exons shorter than 241 nt constitute a majority in all the genomes examined (fig. 1). Interestingly, we found that the longer internal exons are, the more frequently they encode IDRs (predicted by DISOPRED3 (Jones and Cozzetto 2015)) in all species (fig. 1). The correlation coefficients are all positive and significantly different from 0 (supplementary table S1, *P*<0.001, *t*-test, Supplementary Material on line). To assess if the observation is robust to IDR prediction methods used, we used two other programs: DICHOT (Fukuchi et al. 2011) and POODLE-L (Hirose et al. 2007). We found a positive correlation without exception irrespective of IDR prediction methods (supplementary figs. S1 and S2, Supplementary Material on line). The correlation coefficients are all significantly different from zero (*t*-test, *P*<0.001) if DISOPRED3 assignments of IDRs are used and the same goes true for 13 out of 16 cases if DICHOT or POODLE-L is employed for IDR prediction (table S1, Supplementary Material on line). We also examined if long internal exons have high cytidine contents as reported on human exons (Kawachi et al. 2021). Though longer internal exons of most species displayed a tendency to have higher cytidine contents, those of *A. thaliana* did not (supplementary fig. S3, Supplementary Material on line). The inclusion of long internal exons is thus unlikely to be universally mediated by the SRSF3 splicing factor.

**Fig. 1.**
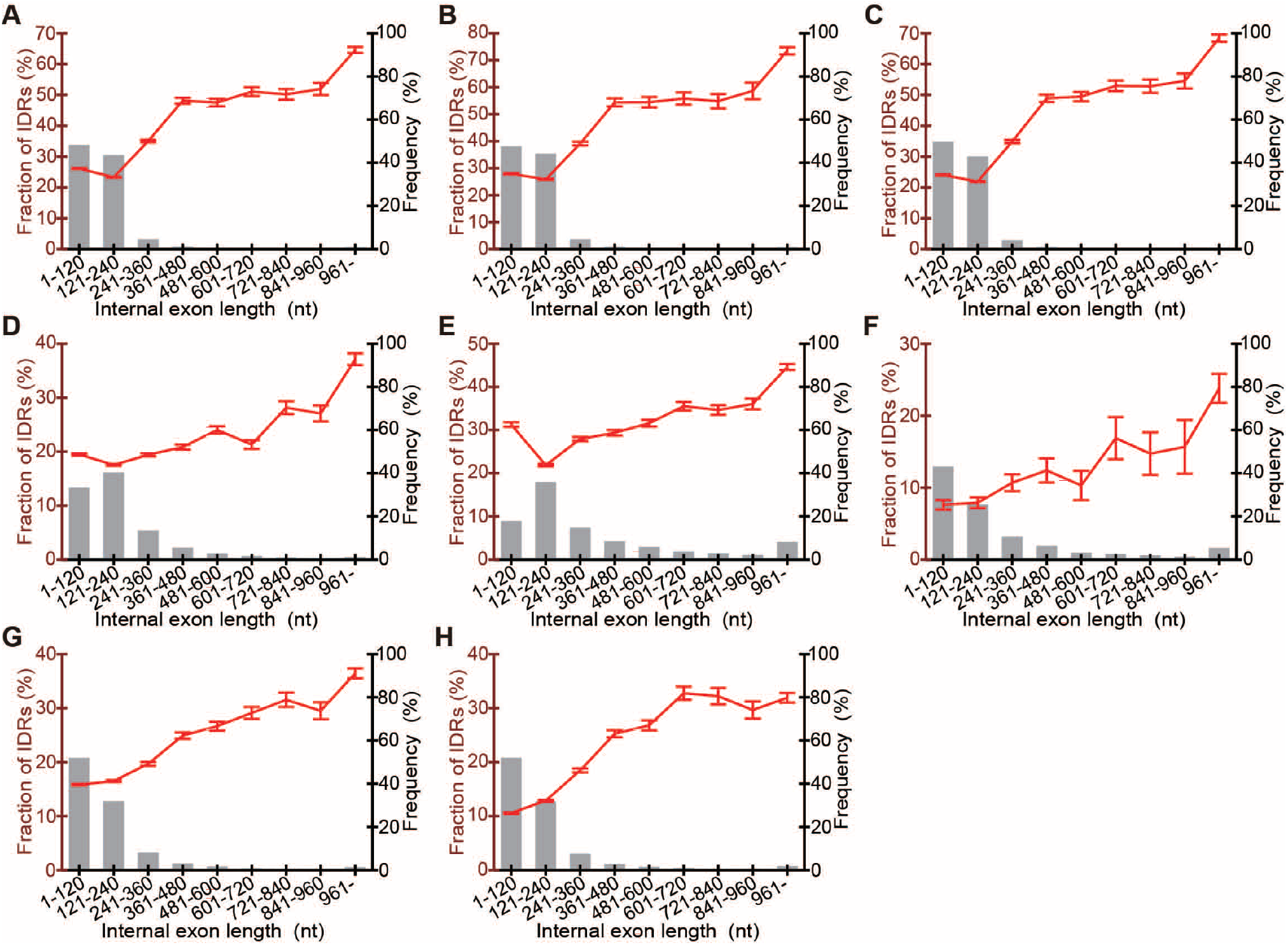
Long internal exons constitute a minority and tend to encode IDRs. The frequencies of occurrence of internal exons in the indicated length ranges are shown by bar graphs (right axis), while the mean fractions of IDRs encoded by the internal exons in each length range together with standard errors of the mean (SEMs) are displayed by line graphs (left axis). The species presented are (*A*) *H. sapiens*, (*B*) *M. musculus*, (*C*) *R. norvegicus*, (*D*) *C. elegans*, (*E*) *D. melanogaster*, (*F*) *S. pombe*, (*G*) *A. thaliana*, and (*H*) *O. sativa*.

### Genes with long internal exons preferentially have nucleus-associated functions

To identify functions of genes with long internal exons, we analysed gene ontology (GO) terms that are enriched in such genes in *H. sapiens, M. musculus, S. pombe*, and *A. thaliana* as these are the only ones in the eight species for which a majority of proteins have been annotated in SwissProt/UniProt (UniProt Consortium 2021), a high-quality protein database. The genes with internal exons longer than 720 nt are annotated with the following GO terms significantly more often (chi-square test, *P*<0.05, the number of genes in each range is shown in supplementary table S2, Supplementary Material on line) in all the species examined: nucleus (GO:0005634), nucleoplasm (GO:0005654), nuclear body (GO:0016604), mRNA processing (GO:0006397), double-strand break repair (GO:0006302), and microtubule (GO:0005874). The same analysis of the genes with extremely long exons (>960 nt) identified four GO terms that are significantly highly represented (chi-square test, *P*<0.05): nucleoplasm, nuclear body, mismatch repair complex (GO:0032300), and microtubule. Among the identified GO terms in both analyses, all except for microtubule are nucleus-specific functions. Besides their roles in intracellular transport and cytoskeleton organisation, microtubules drive chromosome movements and induce nuclear envelope breakdown during cell division. Thus, genes with long internal exons preferentially have nucleus-associated functions.

### The small bang model

One explanation for the tendency of long internal exons to encode IDRs is that some short internal exons that existed in early eukaryotes elongated and most of the elongated sections encoded IDRs. Intron sliding (Rogozin et al. 2012), exonisation of introns, and repeat sequence expansion are possible mechanisms of internal exon elongation. What kind of primordial short exons expanded in length? An internal exon that encodes a portion of an SD (stippled in fig. 2(*A*)), which we call a constrained exon, is presumably resistant to elongation because the inserted segment(s) inside the SD would be structurally destabilizing in most cases and thus would be rarely tolerated. We thus considered it probable that internal exons that expanded were mostly unconstrained exons. Moreover, we thought it likely that the added sections encoding IDRs facilitated recombination to give rise to genes encoding multiple SDs: IDRs are likely to promote domain accretion as they serve to keep a proper spacing between SDs, which is crucial in many protein functions (Sturm and Herr 1988; van Leeuwen et al. 1997; Brodsky et al. 2020).

**Fig. 2.**
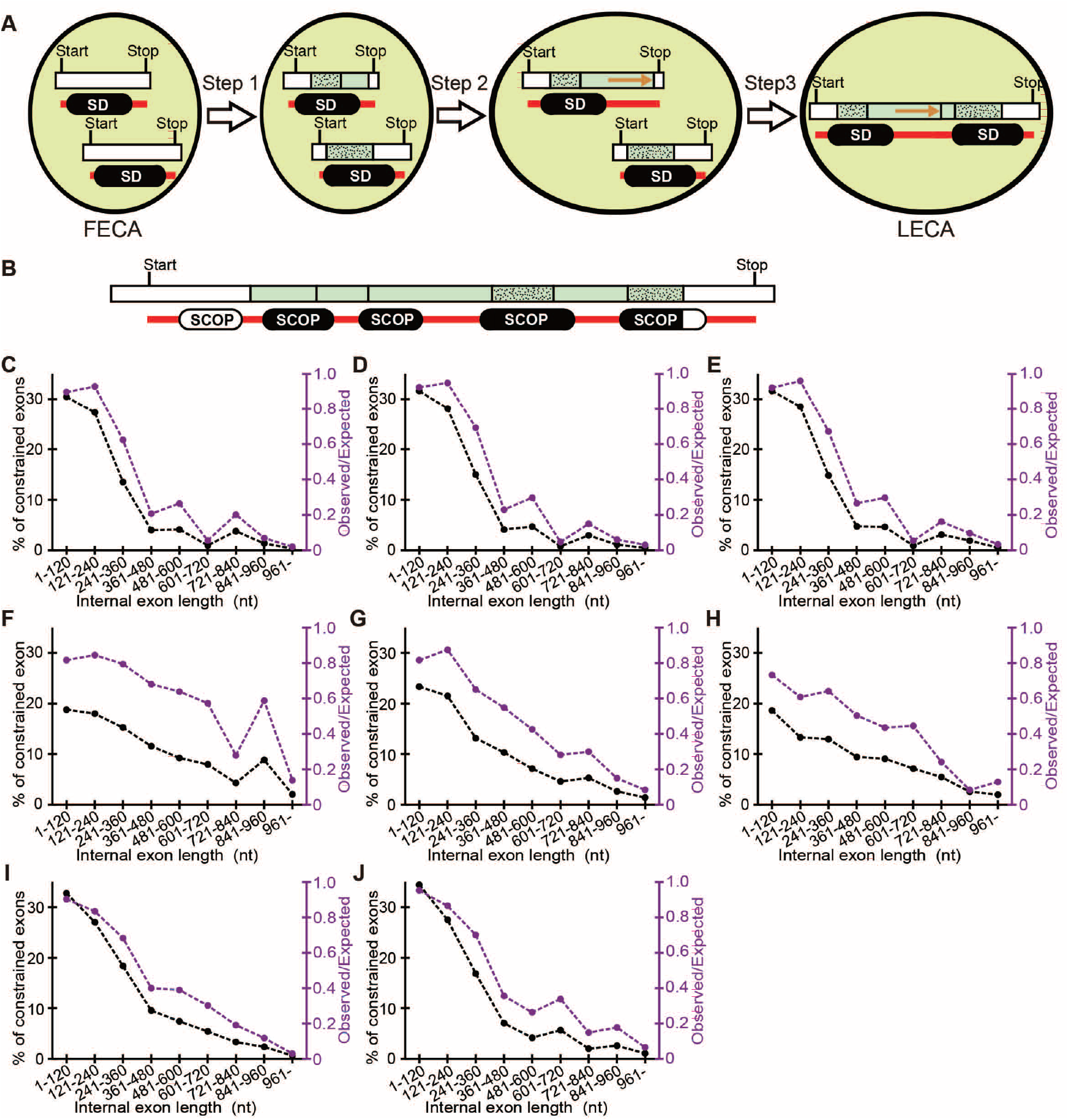
The small bang model and the tendency of longer internal exons to contain less constrained exons. (*A*) The small bang model. The terminal and internal exons are shown as white and pale green rectangles, respectively. Internal exons both of whose boundaries fall inside a SCOP domain are classified as constrained exons (stippled). The IDRs and SDs in the encoded proteins are respectively presented as red bars and rounded black rectangles. The orange arrows indicate an elongated section. (*B*) A hypothetical case presented as in (*A*) except that SDs are replaced by SCOP domains. Only the sections of SCOP domains encoded by the internal exons (black) are taken into account. (*C-J*) The observed fraction of constrained exons (black, left axis) and the ratio of the observed fraction to the expected fraction (purple, right axis) are plotted. The species are (*C*) *H. sapiens*, (*D*) *M. musculus*, (*E*) *R. norvegicus*, (*F*) *C. elegans*, (*G*) *D. melanogaster*, (*H*) *S. pombe*, (*I*) *A. thaliana*, and (*J*) *O. sativa*.

We therefore propose the small bang model from the proteomic viewpoint (fig. 2(*A*)): the genes of the FECA had no introns and most of them encoded a single SD, just like prokaryotic genes. Most genes subsequently divided into exons (step 1). Then a plethora of unconstrained exons expanded with the addition of segments that encode IDRs (step 2). The acquired IDRs provided new beneficial functions to the proteins such as efficient binding of transcribed DNA sequences and facilitation of condensation, most often in the nucleus. Some of the genes with extended exons recombined with each other to give rise to those encoding multiple SDs with the IDRs in the extended segment serving as a spacer between SDs (step 3). As they conferred a selective advantage, these events occurred multiple times to produce the LECA. According to this model, the LECA proteome was comprised mostly of proteins having multiple SDs and a significant fraction of IDRs, accounting for the dichotomy between prokaryotic and eukaryotic proteins.

### Long internal exons are mostly unconstrained exons

Our model would predict that long internal exons in extant eukaryotes are mostly unconstrained exons, that is, the longer internal exons are, the smaller the observed fraction of constrained exons will be. To test this prediction, we used SCOP domains (Andreeva et al. 2019) in our analysis (fig. 2(*B*)), because SD boundaries are not easily identifiable from the outputs of IDR prediction programs as one SD is often artifactually subdivided into multiple segments. Notice that the absence of a SCOP domain alignment does not automatically mean that the segment is an IDR, because SDs whose homologues have not been structurally determined remain unclassified by SCOP and thus may be latent in the segment. The results (fig. 2(*C*)-(*J*)) verify the prediction: in all species examined, both the fraction of constrained exons and the observed to expected ratio decrease as internal exons become longer. The correlation coefficients are all negative and significantly different from zero (supplementary table S3, *n* = 9, *P*<0.001, *t*-test, Supplementary Material on line). In general, expected fractions of constrained exons in long internal exons are low (supplementary table S3, Supplementary Material on line) since SCOP domains longer than the region encoded by long exons are rare (Materials and Methods). Presumably “latent” SCOP domains, i.e., those that have not been identified, would increase both the observed and expected fractions to the same extent, and would essentially not affect our results.

### Insertions preferentially occur in unconstrained exons and mostly encode IDRs

Is it possible to identify elongated segments and determine if they preferentially occur in unconstrained exons and encode mostly IDRs? Since unequivocal identification of such cases that had happened before the emergence of the LECA is difficult, we searched for cases of internal exon elongation after the rat-mouse divergence by comparison of human, mouse, and rat orthologues. (The generally high reliability of human, rat, and mouse data prompted us to analyse the three species). We identified the segments in internal exons that exist only in the rat or mouse genome and regarded them as those inserted after the divergence: the mouse/rat-specific segment is unlikely to have been deleted twice in the human lineage and in the rat/mouse lineage. To obtain segments inserted within internal exons (green in fig. 3(*B*) and sky blue in fig. 3(*F*)), we removed inserted segments that coincide with entire exons (gray in fig. 3(*A*) and (*E*)). Although this method cannot capture all the cases of internal exons elongated after the rat-mouse divergence, the selected cases are probably genuine ones.

**Fig. 3.**
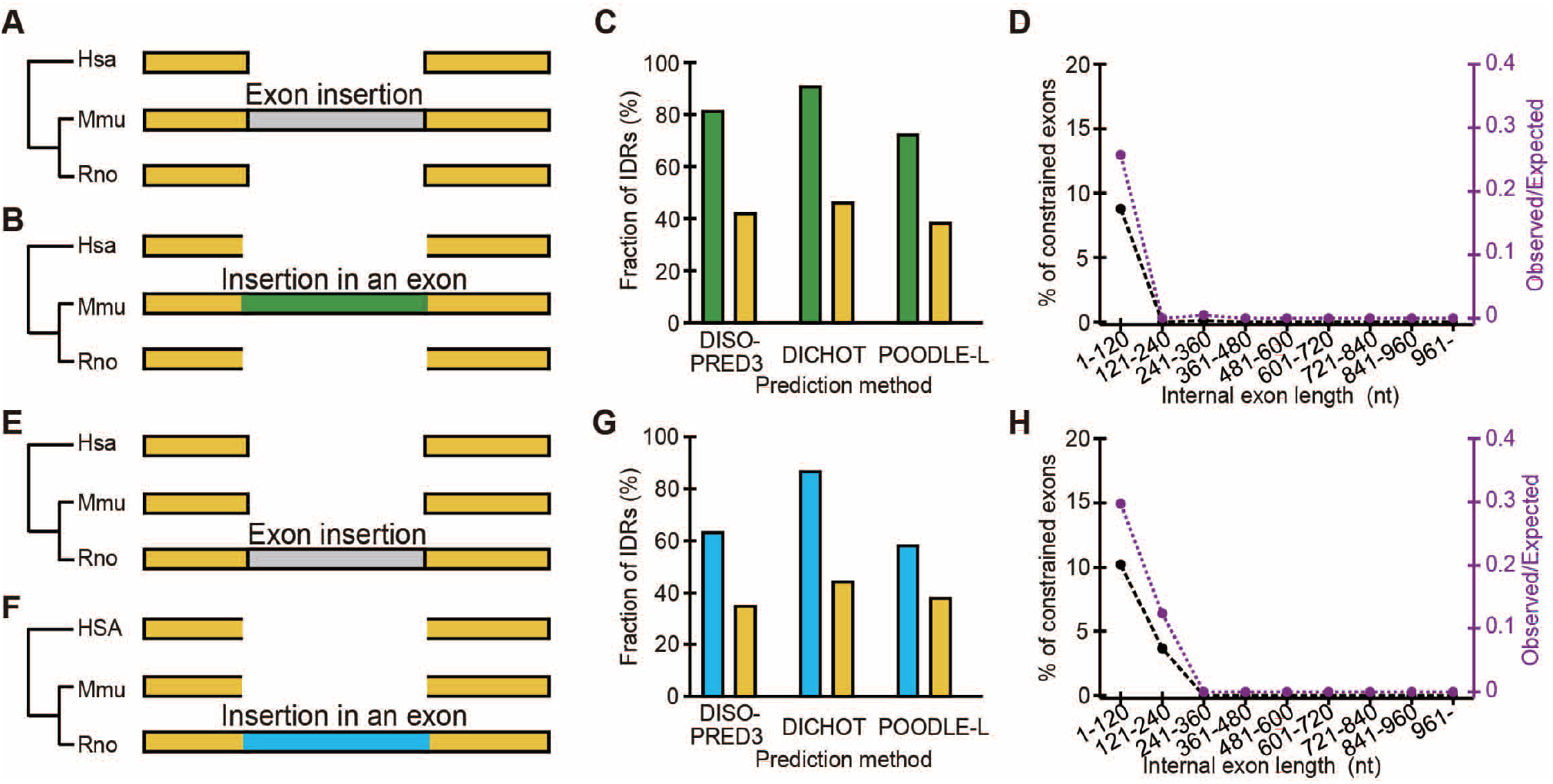
Segments inserted in internal exons after the rat-mouse divergence encode mostly IDRs and preferentially occur in unconstrained exons. (*A-D*) Mouse-specific insertions. (*E-H*) Rat-specific insertions. (*A, B*, *E*, and *F*) The rectangles represent internal exons. (*C* and *G*) The fraction of IDRs in segments inserted in internal exons (green and sky blue, respectively) and that in the rest of internal exons of all genes (yellow) by different prediction methods. (*D* and *H*) The observed fraction of constrained exons in inserted segments (black line, left axis) and the ratio of the observed fraction to the expected fraction (purple, right axis) in each length range of internal exons. The expected fractions are those of all internal exons.

A majority of insertions turned out to be in short (1- 360 nt) internal exons (supplementary table S4, Supplementary Material on line). The segments inserted in internal exons encode mostly IDRs with the fractions much higher than those of the rest of internal exons, irrespective of the prediction methods used (fig. 3(*C*) and (*G*)). We classified internal exons with insertions according to exon length ranges and calculated the average fraction of constrained exons in each length range; due to the preponderance of unconstrained exons in long internal exon ranges (fig. 2(*C*)-(*J*)), we thought it likely that insertions in longer internal exons occur predominantly in unconstrained exons. The fractions of constrained exons are extremely low and are in all cases less than or equal to the expected values (fig. 3(*D*) and (*H*), supplementary table S3, Supplementary Material on line). As the fractions are nearly zero in longer exon length ranges, insertions in longer exons are mostly confined to unconstrained exons.

Four examples of segments inserted in internal segments are presented (fig. 4). The first one (fig. 4(*A*)) is a case in which an internal exon likely expanded by sequence repetition in the rat lineage, adding an IDR-encoding segment as previously suggested (Tompa 2003). The next one (fig. 4(*B*)) represents an example in which the 3’ splicing site of an internal exon was presumably altered in the mouse lineage to incorporate a previously intronic segment in the exon. An alteration of the 5’ splicing site of an internal exon accounts for the case shown as fig. 4(*C*) and the inserted segment also encodes an IDR. Two previously contiguous internal exons apparently formed one long exon by exonization of the intervening intron (fig. 4(*D*)) and the inserted segment again encodes an IDR. The latter three are cases of alternative splicing that affected exons that previously existed. The fact that the extended segments encode IDRs dovetails with the report that alternatively spliced fragments are mostly associated with IDRs (Romero et al. 2006). All the presented inserted segments are located in unconstrained exons. These findings are congruent with the notion that most long internal exons were produced by insertion of IDR-encoding segments to unconstrained exons. We note that the length distribution of internal exons is likely to be in equilibrium in extant eukaryotes: insertions are offset by deletions, since otherwise, some internal exons would lengthen indefinitely.

**Fig. 4.**
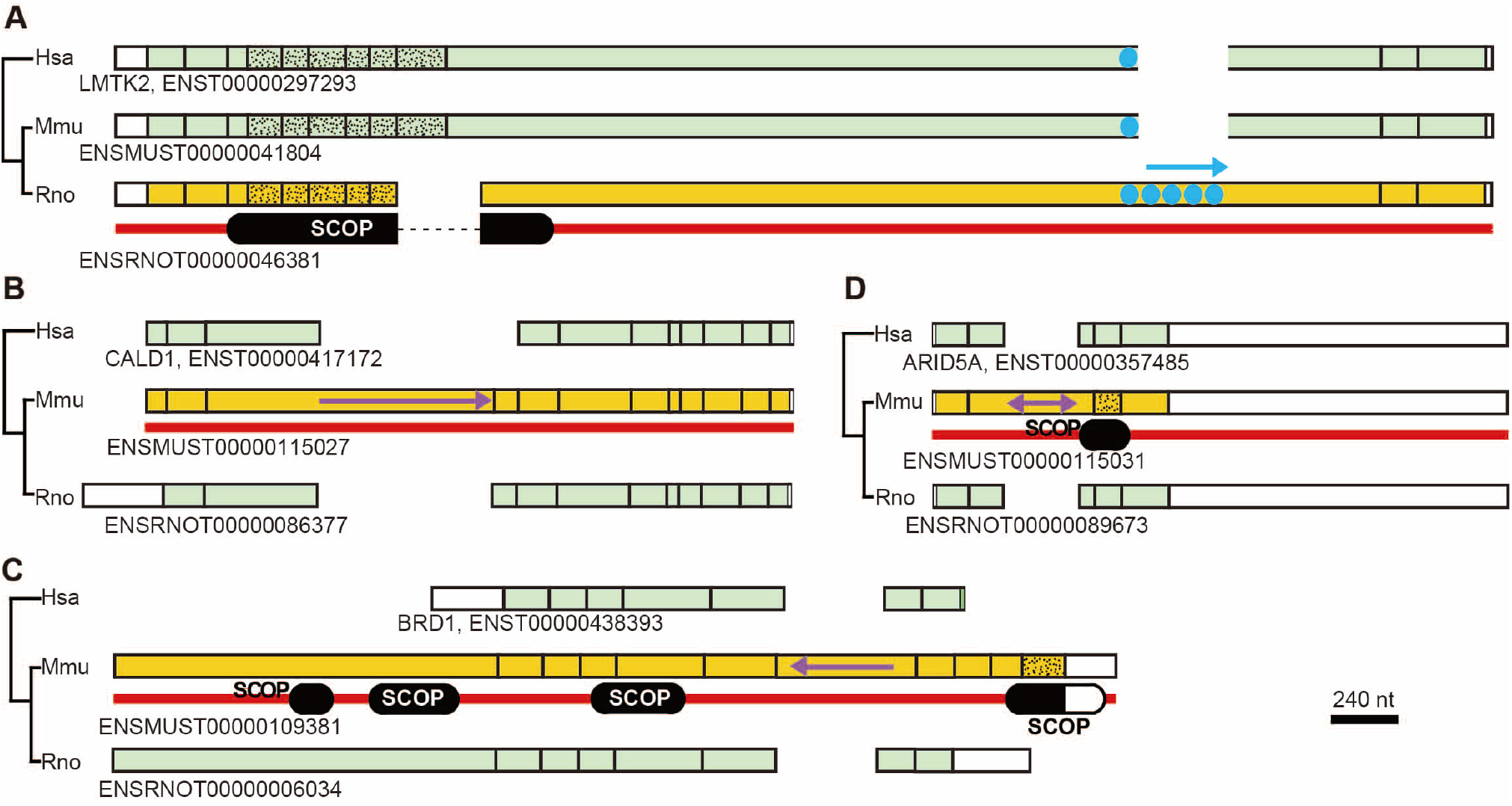
Examples of elongated internal exons. Human, mouse, and rat orthologues are drawn with corresponding exons aligned vertically on the indicated scale. The same phylogenetic tree was added to the left of each pane. The coding regions of the terminal exons (white), those with insertion in internal exons (yellow), and the rest of internal exons (pale green) of orthologues are shown with the constrained exons stippled. The single-headed arrows signify the probable directions of expansion, while the double-headed arrow indicates an exonised intron. The transcript identifications and human gene names are added to the right of the phylogenetic trees and the SCOP domains in encoded proteins are depicted beneath exons. No SCOP domains were assigned in (*B*).

### An SD followed by a long internal exon frequently has another SD downstream

In step 3 of the small bang model, many genes with long internal exons acquired SDs downstream (fig. 2(*A*)). This would predict that genes encoding an SD tend to encode another SD downstream if a long internal exon exists downstream of the exon(s) encoding the first SD. To test this prediction, we identified segments between SCOP domains (group A, indicated by the orange arrow in fig. 5(*A*)) with the expectation to frequently find long internal exons in the segments that facilitated acquisition of the downstream SCOP domains. For comparison, we chose the segment downstream of the last SCOP domain in each gene (group B, the blue arrows in fig. 5(*A*)), as it is predicted not to contain a long internal exon at high frequency as it has not facilitated the addition of SCOP domain(s) downstream. If the model were correct, the length of the longest exon in group A would tend to be longer than that of group B.

**Fig. 5.**
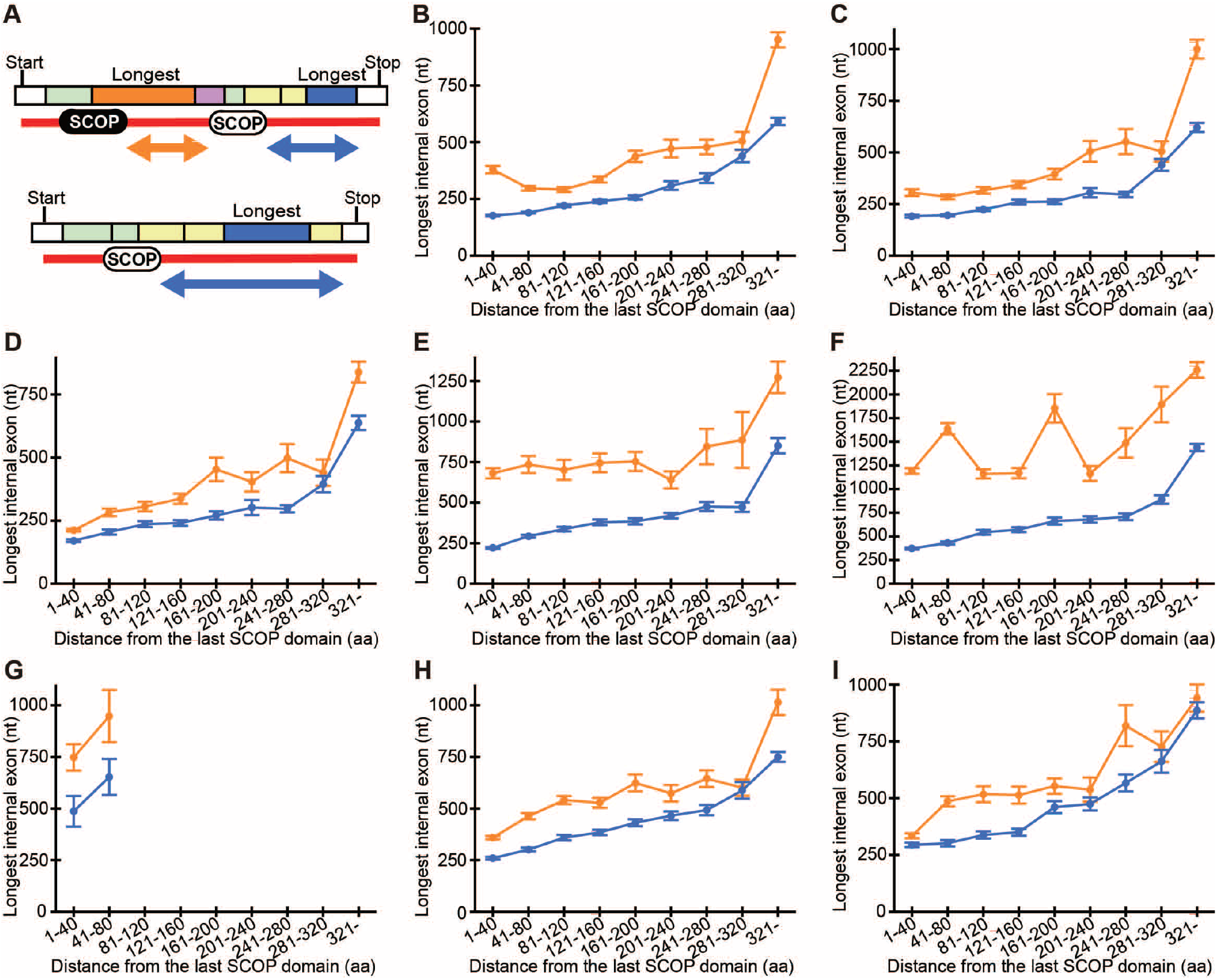
A SCOP domain followed by a long internal exon frequently has another SCOP domain downstream. (*A*) Hypothetical cases to explain the procedure. Internal exons are coloured. If multiple SCOP domains exist, the longest internal exon among the candidates (orange and pink rectangles) in the inter-SCOP region (orange arrow) is identified (group A). In the regions downstream of the last SCOP domain (blue arrows), the longest internal exons among the candidates (yellow and blue rectangles) are determined (group B). (*B-I*) The mean lengths and SEMs of the longest internal exons in group A (orange) and group B (blue) segments are shown as solid lines. The species plotted are (*B*) *H. sapiens*, (*C*) *M. musculus*, (*D*) *R. norvegicus, (E) C. elegans, (F) D. melanogaster*, (*G*) *S. pombe*, (*H*) *A. thaliana*, and (*I) O. sativa*.

We determined the longest exon in each group A segments in genes with SCOP domain(s) and calculated the mean length in each range of segment length (orange in fig. 5 (*B*)-(*I*)). We required both groups A and B to have at least 25 samples for the mean to be calculated (“minimum number requirement”, supplementary table S5, Supplementary Material on line). We also identified the longest exon in the group B segments in all genes with SCOP domain(s) and computed the mean length in each bin of distance from the last SCOP domain (blue in fig. 5 (*B*)-(*I*)). Without exception, in all the eight species examined, the longest internal exons tend to be longer if a SCOP domain is followed by another SCOP domain. This result comports with our model. One concern is that identification of “latent” SCOP domains may change the result. To assess the effect of latent SCOP domains, we randomly selected half of SCOP domains and carried out the same analyses with the selected SCOP domains. Although the number of samples was reduced (supplementary table S6, Supplementary Material on line), the results are essentially the same as before (supplementary fig. S4, Supplementary Material on line). (No mean values were calculated for *S. pombe* as the sample numbers in all the distance ranges failed to satisfy the minimum number requirement (supplementary table S6, Supplementary Material on line)). Thus, the identification of latent SCOP domains would probably not affect the conclusion.

## Discussion

### Three lines of evidence support the small bang model

The small bang model accounts for the important dichotomy between prokaryotic and eukaryotic proteins: eukaryotic proteins have a higher fraction of IDRs and more SDs than prokaryotic counterparts. Firstly, the tendency of longer internal exons to encode more IDRs supports this model. Although the correlation coefficients are positive and are significantly different from zero in all species if IDRs are predicted by DISOPORD3, they are small: 0.098-0.152 (fig. 1 and supplementary table S1, Supplementary Material on line). We also note that the slopes in some species, especially in *S. pombe* (fig. 1(*F*)), are small. We consider it likely that massive exon losses that had occurred in some lineages of eukaryotes after the LECA weakened the initial signal. The fact that the trend is detectable in all the eight species, however, makes it plausible that the tendency is traceable to the LECA. The same signal is predicted to be detectable in all other eukaryotes with a significant number of internal exons.

Genes with long internal exons are enriched with those with nucleus-associated functions. Considering the finding that long internal exons preferentially encode IDRs, this observation comports with the report that IDRs play crucial roles in transcription factors (Minezaki et al. 2006). Some of the newly acquired IDRs may have participated in the formation of biomolecular condensates that regulate multitudes of cellular functions such as transcription, conferring an evolutionary advantage.

Secondly, the model is consistent with the observed low frequency of constrained exons in long internal exons in all the eight eukaryotes analysed. It is likely that many unconstrained exons elongated, with the elongated sections encoding mostly IDRs. If the model were correct, this phenomenon would be observed in other eukaryotes with a large enough number of internal exons. The identified cases of internal exons that probably elongated in the mouse or rat genome are compatible with our model.

Lastly, we found in all the eight eukaryotes examined that genes encoding an SD tend to have another SD downstream if a long internal exon exists in the downstream section. This result is also in agreement with the model. According to the model, this tendency must be universally observed in eukaryotes that retain a large enough number of internal exons. The elongated sections of internal exons that mostly encoded IDRs probably facilitated recombination to add SDs. We acknowledge that this piece of evidence is also compatible with the alternative model with the ordering of events reversed, namely, domain accretion (step 3 in fig. 2(*A*)) occurred prior to the expansion of unconstrained exons (step 2 in fig. 2(*A*)). However, we consider our proposed model more plausible: exon elongation mostly produces segments encoding IDRs, which are likely to enable recombination that adds SDs with proper spacing. As many multidomain proteins provide novel functions, this step plausibly conferred a selective advantage. Since recombination events in the alternative model are predicted to produce dysfunctional genes encoding juxtaposed SDs as well as disrupted SDs, it is less plausible than our model.

### Significance and predictions of the small bang model

Division of genes into exons (step 1 in fig. 2(*A*)) necessitates nucleus-cytosol compartmentalization because mRNA splicing must be completed before translation starts (Martin and Koonin 2006). Thus, if the small bang model were valid, steps 2 and 3 would be placed after the evolution of the nucleus. Exon expansion that added IDRs to the encoded proteins (step 2) enabled transcription factors to efficiently and precisely control gene expression of a large eukaryotic genome (Minezaki et al. 2006). It also drove the formation of the droplet state under appropriate conditions that facilitate various cellular functions including transcription and DNA repair (Fuxreiter and Vendruscolo 2021). We propose that the elongated exons encoding IDRs promoted recombination to produce functional genes encoding multidomain proteins (step 3) that substantially increased the functional complexity of the LECA (Koonin 2000; Koonin et al. 2002; Tordai et al. 2005).

Having acquired IDRs and multidomain proteins necessary for efficient cellular functions, many lineages of eukaryotes such as most unicellular eukaryotes experienced a massive loss of introns after the LECA, as they no longer gave a significant selective advantage. The putative evolutionary advantage provided by introns through attenuation of R-loop formation (Niu 2007) is probably minimal as introns have been entirely lost in a number of extant eukaryotes. On the other hand, the existence of introns in early eukaryotes made it possible for many unconstrained exons to expand with the added regions encoding mostly IDRs, which in turn facilitated domain accretion. The small bang model proposes that eukaryotes that acquired introns gained a selective advantage through acquisition of IDRs and thus augments the exon theory of genes. Moreover, the IDRs encoded by the elongated sections of unconstrained exons decrease the frequency that the boundaries of exons align with the boundaries of structural domains of the encoded proteins. Our model thus explains the reported low frequency of their alignment (Liu and Grigoriev 2004, Smithers et al. 2019).

We note that a few lineages of eukaryotes such as metazoans apparently experienced intron gain (Csuros et al. 2011), possibly producing exons exclusively encoding IDRs. Although this event is expected to weaken the signals we analyzed, the universal presence of signals in the eight species examined supports our model. A number of exons encoding IDRs only may have emerged before the LECA and greatly facilitated exon shuffling. However, we find it hard to test this possibility since the appearance of such exons prior to the LECA is nearly indistinguishable from their emergence posterior to the LECA.

In addition to the aforementioned universality of reported results, the model predicts that the lengths of constrained exons are more evolutionarily conserved than those of unconstrained exons. Another prediction is that genes with longer internal exons have more novel combinations of SDs that do not exist in prokaryotes. Future research will hopefully test and further refine the model.

## Materials and Methods

All the sequence, exon and orthologue data used in this study were downloaded from the Ensembl database (Cunningham et al. 2022). Though we analysed all variants, we made the exons non-redundant except for the identification of the longest internal exon by in-house C programs to avoid double counting. IDRs were predicted by DISOPRED3 (Jones and Cozzetto 2015), DICHOT (Fukuchi et al. 2011), and POODLE-L (Hirose et al. 2007), while SCOP domains (Andreeva et al. 2019) were assigned in an intermediate step in DICHOT prediction.

An internal exon is considered constrained if it encodes a protein region that is entirely contained in one SCOP domain. (Specific examples of constrained exons are provided in fig. 4 (stippled)). Using in-house C scripts, we analysed the proteins to calculate the fraction of internal exons encoding SCOP domains to get the expected occurrence of constrained exons. If L is the total length encoded by internal exons rounded to the nearest integer, the probability of an exon encoding y amino acids to have both boundaries located inside of a SCOP domain of length x_i_ is given by (x_i_– y–1)/(L-y+1) if x_i_ > y+1 and is zero otherwise. The total probability is the sum of the probability over all SCOP domains encoded by internal exons.

Mouse or rat specific insertion segments were identified from the MAFFT (Katoh and Standley 2013) alignments of orthologous protein sequences with default settings. We wrote C programs to assess enrichment of GO terms according to SwissProt/UniProt (UniProt Consortium 2021) annotations (release 2021_04).

In-house C scripts identified the longest internal exon among candidate exons and calculated the mean length and SEM in each range of distance from the last SCOP domain. In the programs, an internal exon is considered as a candidate of the longest internal exon of group A if the exon encodes the C-terminal end of the last SCOP domain. In group A, an internal exon is also considered as a candidate if the beginning of the exon encodes a residue between contiguous SCOP domains. An internal exon is included as a candidate of the longest internal exon of group B if the last residue the exon encodes is downstream of the last SCOP domain. Genes encoding no SCOP domains are neglected in both groups.

## Supplementary Material

Supplementary data are available at *Molecular Biology and Evolution* on line.

## Acknowledgments

We thank K. Hosoda for setting up a system to run POODLE-L in house and W-C. Goh for a critical reading.

## Data Availability

The custom scripts used in this study (supplementary table S7, Supplementary Material on line) are available in a GitHub repository (https://github.com/hommadekkai/evolution-leca.git).

