## SupplementalMaterials for "Exon Elongation Added Intrinsically Disordered Regions to the Encoded Proteins and Facilitated the Emergence of the Last Eukaryotic Common Ancestor"

4 supplementary figures and 7 supplementary tables

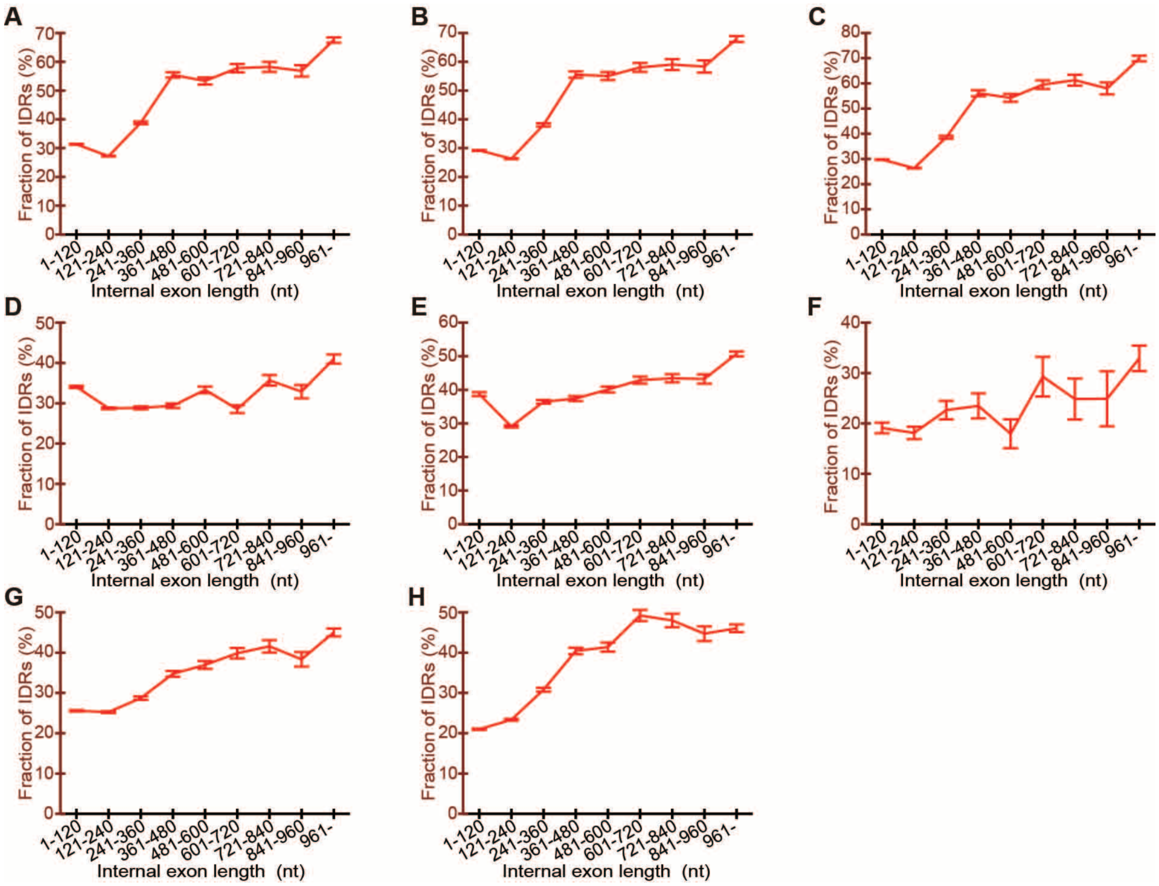


**Supplementary Fig. S1. DICHOT results show that long internal exons tend to encode IDRs.** The mean fractions of IDRs encoded by the internal exons in the length rages with SEMs are displayed by line graphs. The IDRs were predicted by DICHOT and are presented as in Fig. 1.

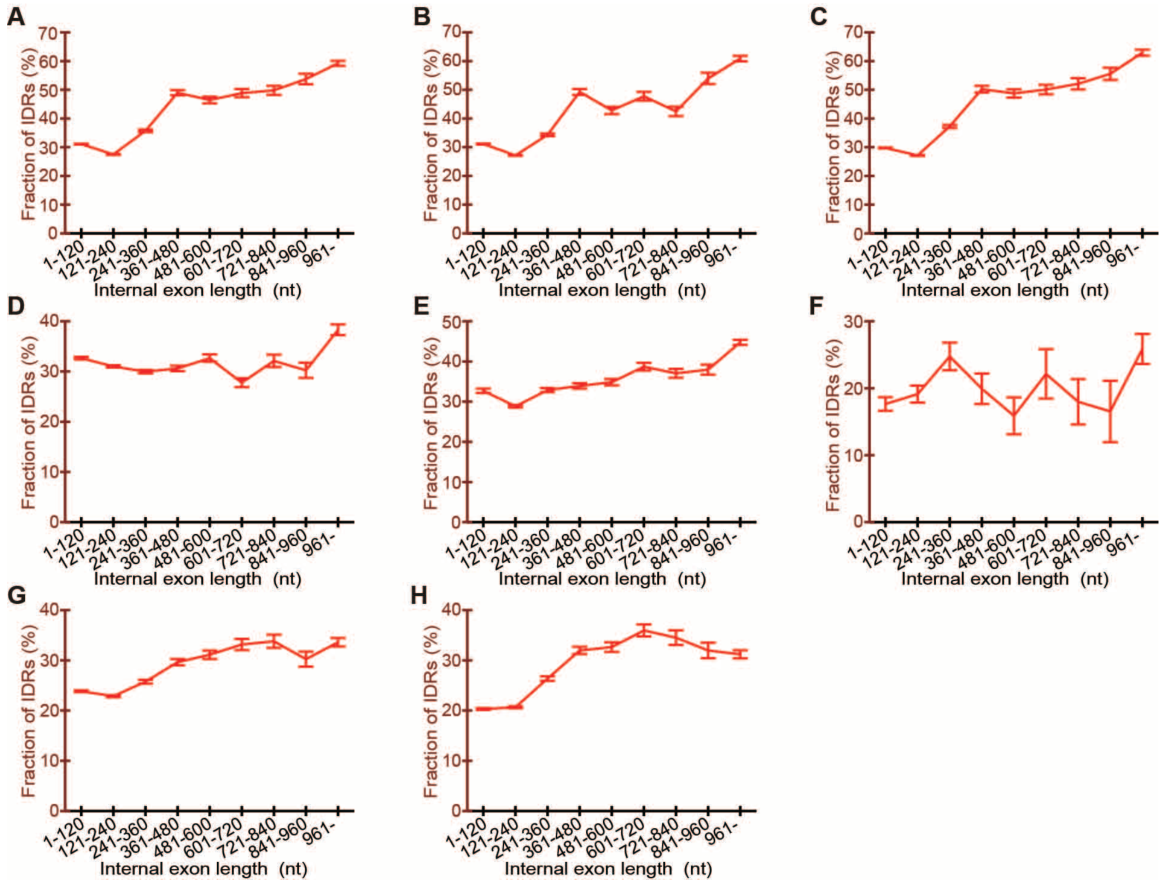


**Supplementary Fig. S2. POODLE-L results show that long internal exons tend to encode IDRs.** The mean fractions of IDRs encoded by the internal exons in the length rages with SEMs are displayed by line graphs. The IDRs were predicted by POODLE-L and are shown as in Fig. 1.

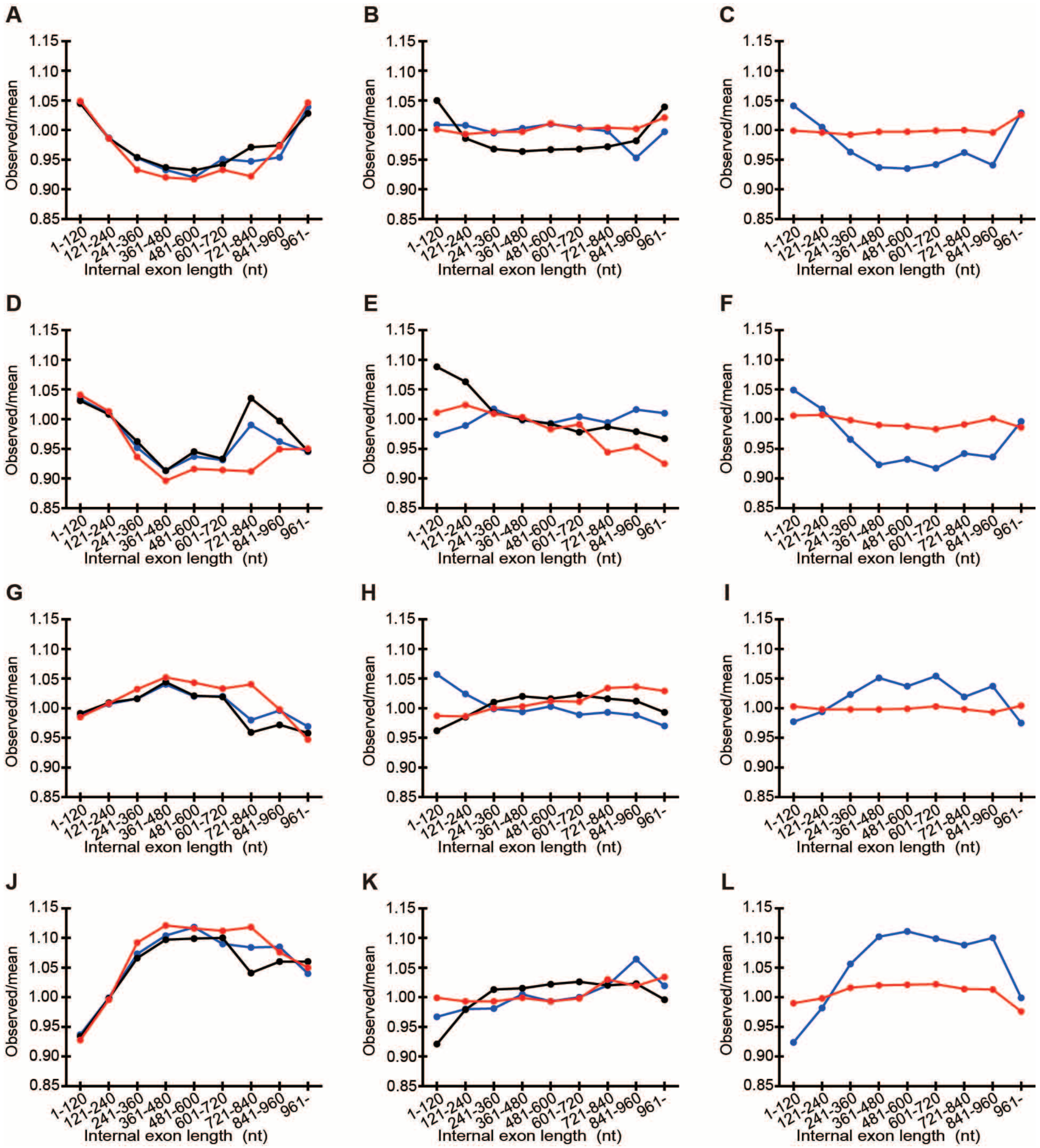


**Supplementary Fig. S3. Long internal exons in most but not all species have high cytidine contents.** The observed frequency of a nucleotide in the internal exons in each length bin is divided by the average frequency of the nucleotide in all the internal exons in the species. The ratios in each species coloured are shown as line graphs. The nucleotides plotted are *(A-C)* adenine, (*D-F*) thymidine, (*G-I*) guanosine, and (*J-L*) cytidine. (*A*, *D*, *G*, and *J*) *H. sapiens* (red), *M. musculus* (black), and *R. norvegicus* (blue), (*B*, *E*, *H*, and *K*) *C. elegans* (red), *D. melanogaster* (black), and *S. pombe* (blue), (*C*, *F*, *I*, and *L*) *A. thaliana* (red) and *O. sativa* (blue).

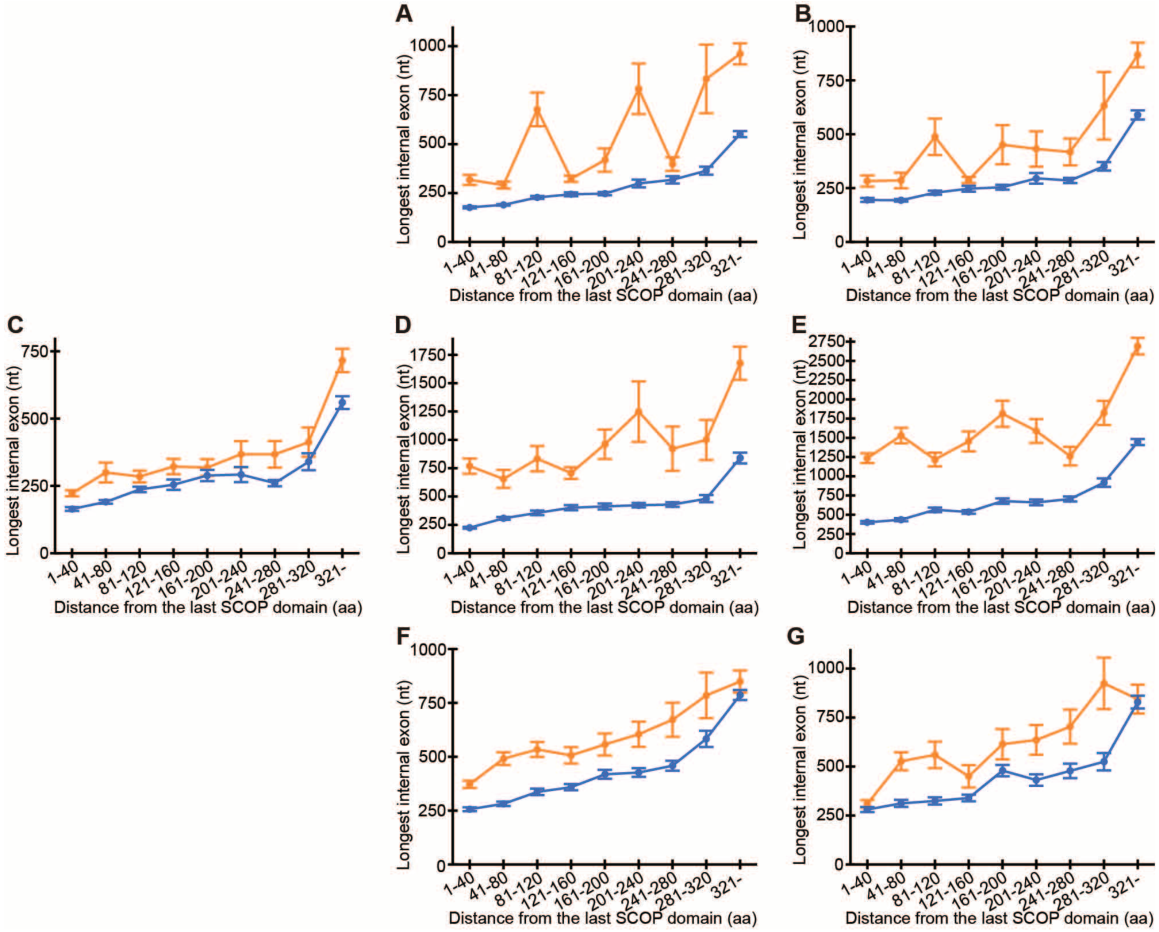


**Supplementary Fig. S4.** A SCOP domain followed by a long internal exon frequently has another SCOP domain downstream even if only half of SCOP domains are randomly selected and used for analysis. The mean lengths and SEMs of the longest internal exons in group A (orange) and B (blue) segments are shown as solid lines if the numbers of both groups are in excess of 20. The species plotted are (*A*) *H. sapiens*, (*B*) *M. musculus*, (*C*) *R. norvegicus*, (*D*) *C. elegans*, (*E*) *D. melanogaster*, (*F*) *A. thaliana*, and (*G*) *O. sativa*.

**Supplementary Table S1.** The number of samples and correlation coefficients (*R*s) between internal exon length and per cent IDRs.

| Species | Hsa | Mmu | Rno | Cel | Dme | Spo | Ath | Osa |
|---|---|---|---|---|---|---|---|---|
| Sample number | 187,484 | 178,114 | 166,917 | 87,985 | 37,952 | 2,785 | 100,776 | 85,468 |
| *R* by DISOPRED3 | 0.098 | 0.109 | 0.113 | 0.065 | 0.130 | 0.146 | 0.099 | 0.152 |
| *R* by DICHOT | 0.074 | 0.087 | 0.086 | 0.007* | 0.093 | 0.073 | 0.071 | 0.100 |
| *R* by POODLE-L | 0.066 | 0.065 | 0.077 | 0.009* | 0.097 | 0.020* | 0.048 | 0.074 |

*Not significantly different from 0 at $P<0.001$, while the rest are significantly different from 0 at $P<0.001$.

**Supplementary Table S2.** The number of genes with maximum internal exon lengths in each range.

| Maximum internal exon length (nt) | Hsa | Mmu | Rno | Cel | Dme | Spo | Ath | Osa |
|---|---|---|---|---|---|---|---|---|
| 1-120 | 3,655 | 1,474 | 858 | 616 | 712 | 365 | 2,982 | 2,448 |
| 121-240 | 11,665 | 9,581 | 7,163 | 3,287 | 2,140 | 345 | 7,184 | 7,141 |
| 241-360 | 3,948 | 3,364 | 2,427 | 3,174 | 1,735 | 169 | 4,112 | 3,734 |
| 361-480 | 1,461 | 1,104 | 826 | 2,030 | 1,508 | 130 | 2,058 | 1,796 |
| 481-600 | 941 | 724 | 520 | 1,207 | 1,279 | 66 | 1,323 | 1,039 |
| 601-720 | 674 | 578 | 417 | 889 | 975 | 58 | 800 | 716 |
| 721-840 | 501 | 423 | 287 | 499 | 773 | 53 | 550 | 506 |
| 841-960 | 369 | 323 | 228 | 314 | 656 | 36 | 430 | 439 |
| 961- | 1,600 | 1,354 | 885 | 731 | 2,894 | 145 | 1,374 | 1,541 |

**Supplementary Table S3.** The expected fractions (%) of constrained exons and Spearman's correlation coefficients ($r_s$'s).

| Internal exon length (nt) | HSA | MMU | RNO | CEL | EME | SPO | ATH | OSA |
|---|---|---|---|---|---|---|---|---|
| 1-120 | 33.96 | 34.20 | 34.29 | 23.00 | 28.59 | 25.42 | 36.29 | 36.19 |
| 121-240 | 29.39 | 29.63 | 29.69 | 21.25 | 24.53 | 21.92 | 32.44 | 31.81 |
| 241-360 | 21.63 | 21.61 | 22.07 | 19.15 | 20.21 | 20.21 | 26.87 | 24.05 |
| 361-480 | 19.36 | 17.97 | 17.79 | 16.99 | 18.86 | 18.74 | 23.89 | 19.81 |
| 481-600 | 15.60 | 15.75 | 15.58 | 14.46 | 16.74 | 20.88 | 19.03 | 15.87 |
| 601-720 | 17.54 | 17.09 | 17.30 | 13.87 | 16.28 | 16.06 | 17.87 | 16.72 |
| 721-840 | 18.82 | 19.72 | 19.38 | 15.30 | 17.63 | 22.65 | 17.35 | 13.50 |
| 841-960 | 19.76 | 18.18 | 19.72 | 15.07 | 17.65 | 30.99 | 20.17 | 14.58 |
| 961- | 16.83 | 16.09 | 15.69 | 14.71 | 16.77 | 15.41 | 21.29 | 16.51 |
| $r_s$ between internal exon length and observed fraction | -0.93 | -0.93 | -0.95 | -0.95 | -0.98 | -1.00 | -1.00 | -0.97 |
| $r_s$ between internal exon length and observed/expected ratio | -0.92 | -0.92 | -0.92 | -0.93 | -0.97 | -0.95 | -1.00 | -0.97 |

**Supplementary Table S4.** The number of inserted segments.

| Internal exon length (nt) | Mmu | Rno |
|---|---|---|
| 1-120 | 6753 | 4426 |
| 121-240 | 1863 | 930 |
| 241-360 | 1809 | 470 |
| 361-480 | 974 | 209 |
| 481-600 | 632 | 177 |
| 601-720 | 225 | 103 |
| 721-840 | 86 | 43 |
| 841-960 | 924 | 50 |
| 961- | 7446 | 537 |

**Supplementary Table S5.** The number of segments in each range of distance from the last SCOP domain.

| Group A | Hsa | Mmu | Rno | Cel | Dme | Spo | Ath | Osa |
|---|---|---|---|---|---|---|---|---|
| 1-40 aa | 9925 | 6013 | 3873 | 1671 | 3433 | 74 | 2622 | 1868 |
| 41-80 aa | 2751 | 1766 | 1108 | 601 | 1206 | 33 | 1045 | 743 |
| 81-120 aa | 1738 | 1068 | 671 | 376 | 702 | 17 | 585 | 303 |
| 121-160 aa | 1135 | 765 | 487 | 214 | 431 | 6 | 357 | 200 |
| 161-200 aa | 763 | 488 | 320 | 143 | 279 | 2 | 190 | 143 |
| 201-240 aa | 549 | 369 | 229 | 86 | 252 | 1 | 198 | 79 |
| 241-280 aa | 430 | 248 | 172 | 107 | 191 | 5 | 144 | 67 |
| 281-320 aa | 333 | 198 | 130 | 82 | 162 | 1 | 109 | 74 |
| 321 aa- | 1679 | 1104 | 689 | 290 | 1048 | 10 | 262 | 162 |

| Group B | Hsa | Mmu | Rno | Cel | Dme | Spo | Ath | Osa |
|---|---|---|---|---|---|---|---|---|
| 1-40 aa | 4025 | 2068 | 1315 | 1054 | 1094 | 65 | 2456 | 2024 |
| 41-80 aa | 2120 | 1204 | 803 | 595 | 608 | 26 | 1044 | 794 |
| 81-120 aa | 1462 | 830 | 564 | 410 | 417 | 16 | 798 | 595 |
| 121-160 aa | 988 | 560 | 408 | 318 | 410 | 15 | 653 | 434 |
| 161-200 aa | 732 | 450 | 279 | 237 | 344 | 10 | 472 | 275 |
| 201-240 aa | 563 | 370 | 229 | 210 | 277 | 9 | 349 | 238 |
| 241-280 aa | 472 | 315 | 217 | 152 | 259 | 8 | 285 | 170 |
| 281-320 aa | 369 | 237 | 154 | 120 | 184 | 8 | 207 | 143 |
| 321 aa- | 2029 | 1346 | 870 | 559 | 1146 | 23 | 799 | 455 |

**Supplementary Table S6.** The number of segments in each range of distance from the last SCOP domain if half of SCOP domains are randomly selected and used for analysis.

| Group A | Hsa | Mmu | Rno | Cel | Dme | Spo | Ath | Osa |
|---|---|---|---|---|---|---|---|---|
| 1-40 aa | 2634 | 1556 | 1053 | 450 | 863 | 18 | 664 | 471 |
| 41-80 aa | 860 | 506 | 349 | 171 | 373 | 6 | 259 | 201 |
| 81-120 aa | 817 | 458 | 274 | 136 | 289 | 9 | 198 | 88 |
| 121-160 aa | 540 | 364 | 200 | 92 | 224 | 5 | 128 | 77 |
| 161-200 aa | 419 | 270 | 151 | 81 | 176 | 2 | 89 | 50 |
| 201-240 aa | 395 | 215 | 129 | 54 | 141 | 1 | 94 | 45 |
| 241-280 aa | 270 | 152 | 100 | 53 | 117 | 2 | 62 | 42 |
| 281-320 aa | 235 | 157 | 90 | 50 | 154 | 1 | 54 | 37 |
| 321 aa- | 1266 | 836 | 495 | 238 | 770 | 9 | 169 | 100 |

| Group B | Hsa | Mmu | Rno | Cel | Dme | Spo | Ath | Osa |
|---|---|---|---|---|---|---|---|---|
| 1-40 aa | 2166 | 1110 | 707 | 515 | 601 | 39 | 1305 | 1117 |
| 41-80 aa | 1275 | 774 | 500 | 323 | 414 | 17 | 637 | 499 |
| 81-120 aa | 1069 | 645 | 372 | 276 | 342 | 14 | 580 | 412 |
| 121-160 aa | 800 | 417 | 348 | 229 | 282 | 15 | 490 | 330 |
| 161-200 aa | 635 | 403 | 256 | 173 | 294 | 8 | 342 | 229 |
| 201-240 aa | 534 | 332 | 246 | 199 | 239 | 6 | 278 | 207 |
| 241-280 aa | 474 | 327 | 215 | 147 | 196 | 5 | 258 | 152 |
| 281-320 aa | 418 | 228 | 161 | 92 | 180 | 7 | 225 | 138 |
| 321 aa- | 2522 | 1655 | 1086 | 563 | 1255 | 24 | 919 | 518 |

**Supplementary Table S7.** List of programs used.

| Species | Hsa | Mmu | Rno | Cel |
|---|---|---|---|---|
| Exons | hallexon3.c | mallexon3.c | nallexon3.c | callexon3.c |
| Non-redundant exons | halluniqexon3.c | malluniqexon3.c | nalluniqexon3.c | calluniqexon3.c |
| Non-redundant internal exons | halluniqexon3A.c | malluniqexon3A.c | nalluniqexon3A.c | calluniqexon3A.c |
| Internal exons with coding ranges | hallexon3M.c | mallexon3M.c | nallexon3M.c | callexon3M.c |
| Non-redundant internal exons with coding ranges | halluniqexon3M.c | malluniqexon3M.c | nalluniqexon3M.c | calluniqexon3M.c |
| Nonredundant exons with classification | halluniqexon4AB.c | malluniqexon4AB.c | nalluniqexon4AB.c | calluniqexon4AB.c |
| ORF list of cDNAs | | mcdnaorflist1.c | ncdnaorflist1.c | ccdnaorflist1.c |
| Abbreviated nonredundant exons with classification | habbcdnalist1.c | mabbcdnalist1.c | nabbcdnalist1.c | cabbcdnalist1.c |
| Protein ID-transcript ID correspondence | | mptcorr1.c | tptcorr1.c | |
| %IDR by DISOPRED3 (fig. 1) | hgintexst6M.c | mgintexst6M.c | ngintexst6M.c | cgintexst6M.c |
| Internal exon length distribution (fig. 1) | hlongexgenes1.c | mlongexgenes1.c | nlongexgenes1.c | clongexgenes1.c |
| SCOP analyses (fig. 2) | hscopconstex3.c | mscopconstex3.c | nscopconstex3.c | cscopconstex3.c |
| MAFFT alignments | | hmrmafft1.c | hmrmafft1.c | |
| Species specific regions | | minsseg1.c | tinsseg1.c | |

| | | | | |
|---|---|---|---|---|
| Insertion in internal exons | | mrinsexon1G.c | trinsexon1G.c | |
| %IDR in insertion by DISOPRED3 (fig.3) | | mrinsidr6G.c | trinsidr6G.c | |
| %IDR in insertion by POODLE-L (fig. 3) | | mrinsidr4G.c | trinsidr4G.c | |
| %IDR in insertion by DICHOT (fig. 3) | | mrinsidr5G.c | trinsidr5G.c | |
| %Constrained exons | | minsconstex3.c | tiinsconstex3.c | |
| Longest internal exon downstream of the last SCOP domain (fig. 5, table S5) | hscopadd2AM.c | mscopadd2AM.c | nscopadd2AM.c | cscopadd2AM.c |
| GO frequency | hlongingo3.c | mlongingo3.c | | |
| GO in genes with long internal exons | commonlongingo3.c | commonlongingo3.c | | |
| GO in genes with extremely long internal eons | commonlongingo4.c | commonlongingo4.c | | |
| fig. S1 data | hintexst5M.c | mintexst5M.c | nintexst5M.c | cintexst5M.c |
| fig. S2 data | hgintexst4M.c | mgintexst4MC.c | ngintexst4M.c | cgintexst4M.c |
| fig. S3 data | hexdnacomp1.c | mexdnacomp1.c | nexdnacomp1.c | cexdnacomp1.c |
| fig. S4 data | hscopadd3AM.c | mscopadd3AM.c | nscopadd3AM.c | cscopadd3AM.c |
| table S1 data | halluniqexon3A.c | malluniqexon3A.c | nalluniqexon3A.c | calluniqexon3A.c |
| table S2 data | hscopnumb2.c | mscopnumb2.c | nscopnumb2.c | cscopnumb2.c |
| table S3 data | hscopconstex3.c | mscopconstex3.c | nscopconstex3.c | cscopconstex3.c |
| table S4 data | | minsconstex3.c | tinsconstex3.c | |

| Species | Dme | Spo | Ath | Osa |
|---|---|---|---|---|
| Exons | dallexon3.c | fallexon3.c | aallexon3.c | rallexon3.c |
| Non-redundant exons | dalluniqexon3.c | falluniqexon3.c | aalluniqexon3.c | ralluniqexon3.c |
| Non-redundant internal exons | dalluniqexon3A.c | falluniqexon3A.c | aalluniqexon3A.c | ralluniqexon3A.c |
| Internal exons with coding ranges | dallexon3M.c | fallexon3M.c | aallexon3M.c | rallexon3M.c |
| Non-redundant internal exons with coding ranges | dalluniqexon3M.c | falluniqexon3M.c | aalluniqexon3M.c | ralluniqexon3M.c |
| Nonredundant exons with classification | dalluniqexon4AB.c | falluniqexon4AB.c | aalluniqexon4AB.c | ralluniqexon4AB.c |
| ORF list of cDNAs | dcdnaorflist1.c | fcdnaorflist1.c | acdnaorflist1.c | rcdnaorflist1.c |
| Abbreviated nonredundant exons with classification | dabbcdnalist1.c | fabbcdnalist1.c | aabbcdnalist1.c | rabbcdnalist1.c |
| %IDR by DISOPRED3 (fig. 1) | dgintexst6M.c | fgintexst6M.c | agintexst6M.c | rgintexst6M.c |
| Internal exon length distribution (fig. 1) | dlongexgenes1.c | flongexgenes1.c | alongexgenes1.c | rlongexgenes1.c |
| SCOP analyses (fig. 2) | dscopconstex3.c | fscopconstex3.c | ascopconstex3.c | rscopconstex3.c |
| Longest internal exon downstream of the last SCOP domain (fig. 5, table S5) | dscopadd2AM.c | fscopadd2AM.c | ascopadd2AM.c | rscopadd2AM.c |
| GO frequency | | flongingo3.c | alongingo3.c | |
| GO in genes with long | | commonlongingo3.c | commonlongingo3.c | |

| | | | | |
|---|---|---|---|---|
| internal exons | | | | |
| GO in genes with extremely long internal eons | | commonlongingo4.c | commonlongingo4.c | |
| fig. S1 data | dintexst5M.c | fintexst5M.c | aintexst5M.c | rintexst5M.c |
| fig. S2 data | dgintexst4M.c | fgintexst4M.c | agintexst4M.c | rgintexst4M.c |
| fig. S3 data | dexdnacomp1.c | fexdnacomp1.c | aexdnacomp1.c | rexdnacomp1.c |
| fig. S4 data | dscopadd3AM.c | fscopadd3AM.c | ascopadd3AM.c | rscopadd3AM.c |
| table S1 data | dalluniqexon3A.c | falluniqexon3A.c | aalluniqexon3A.c | ralluniqexon3A.c |
| table S2 data | dscopnumb2.c | fscopnumb2.c | ascopnumb2.c | rscopnumb2.c |
| table S3 data | dscopconstex3.c | fscopconstex3.c | ascopconstex3.c | rscopconstex3.c |
